# Real-time imaging of optic nerve head collagen microstructure and biomechanics using instant polarized light microscopy

**DOI:** 10.1101/2021.10.03.462955

**Authors:** Po-Yi Lee, Bin Yang, Yi Hua, Susannah Waxman, Ziyi Zhu, Fengting Ji, Ian A Sigal

## Abstract

Current tools lack the temporal or spatial resolution necessary to image many important aspects of the architecture and dynamics of the optic nerve head (ONH). We evaluated the potential of instant polarized light microscopy (IPOL) to overcome these limitations by leveraging the ability to capture collagen fiber orientation and density in a single image. Coronal sections through the ONH of fresh normal sheep eyes were imaged using IPOL while they were stretched using custom uniaxial or biaxial micro-stretch devices. IPOL allows identifying ONH collagen architectural details, such as fiber interweaving and crimp, and has high temporal resolution, limited only by the frame rate of the camera. Local collagen fiber orientations and deformations were quantified using color analysis and image tracking techniques. We quantified stretch-induced collagen uncrimping of lamina cribrosa (LC) and peripapillary sclera (PPS), and changes in LC pore size (area) and shape (convexity and aspect ratio). The simultaneous high spatial and temporal resolutions of IPOL revealed complex ONH biomechanics: i) stretch-induced local deformation of the PPS was nonlinear and nonaffine. ii) under load the crimped collagen fibers in the PPS and LC straightened, without torsion and with only small rotations. iii) stretch-induced LC pore deformation was anisotropic and heterogeneous among pores. Overall, with stretch the pores were became larger, more convex, and more circular. We have demonstrated that IPOL reveals details of collagen morphology and mechanics under dynamic loading previously out of reach. IPOL can detect stretch-induced collagen uncrimping and other elements of the tissue nonlinear mechanical behavior. IPOL showed changes in pore morphology and collagen architecture that will help improve understanding of how LC tissue responds to load.

**Highlights:**

- We demonstrate that instant polarized light microscopy allows visualization and quantification of changes in optic nerve head collagen morphology and architecture under dynamic loading
- We show crimped collagen fibers in the peripapillary sclera and lamina cribrosa straightening under load, without torsion and with only small rotations.
- We show that stretch-induced local deformation of the peripapillary sclera was nonlinear and nonaffine.
- We show that stretch-induced lamina cribrosa pore deformation was anisotropic and heterogeneous among pores.
- Our results show this novel imaging technique could help understand the role of collagen microstructure in eye physiology, aging, and in biomechanics-related diseases, such as glaucoma and myopia.

## 1. Introduction

Collagen fibers are a primary load-bearing component of the optic nerve head (ONH), and thus their organization and behavior under load play a central role in the physiology and pathophysiology of the eye. (Coudrillier et al., 2012; Ethier et al., 2004a) Many imaging techniques have been deployed for measuring and/or visualizing the architecture and biomechanics of ONH collagen. These include confocal, (Kang and Yu, 2015; Masters, 1998) nonlinear, (Behkam et al., 2019; Brown et al., 2007; Sigal et al., 2014a) scanning electron, (Quantock et al., 2015; Quigley and Addicks, 1981; Quigley et al., 1983) and transmission electron (Elkington et al., 1990) microscopies, small-angle light scattering, (Girard et al., 2011; Yan et al., 2011) electronic speckle pattern interferometry, (Bianco et al., 2020; Girard et al., 2009) magnetic resonance imaging, (Ho et al., 2014; Ho et al., 2016) ultrasound, (Ma et al., 2020; Ma et al., 2019; Pavlatos et al., 2018; Pavlatos et al., 2016; Qian et al., 2020) and optical coherence tomography. (Fazio et al., 2018; Midgett et al., 2019) Each of these techniques offers a unique combination of resolution, field of view, penetration depth, speed, and tissue specificity. For instance, nonlinear microscopy has high tissue specificity and spatial resolution, but it has low imaging speed and a small field of view. Thus, studies of ONH biomechanics using nonlinear microscopy have been limited to static or quasi-static conditions. (Sigal et al., 2014a) Electronic speckle pattern interferometry allows real-time imaging, and has excellent resolution to resolve sub-fiber level deformations, but does not discern collagen and has extremely low penetration into tissues. (Bianco et al., 2020) Optical coherence tomography works well in-vivo and has therefore been widely deployed for both research (Sigal et al., 2014b) and clinical (Schuman et al., 2020) work. However, its limitations, primarily in resolution and signal penetration, have precluded its use to quantify local ONH tissue architecture and biomechanics.

Over the past several years, polarized light microscopy (PLM) has been demonstrated to allow visualization and quantification of ONH collagen tissues with micrometer-scale resolution over wide regions. (Brazile et al., 2018; Gogola et al., 2018a; Gogola et al., 2018b; Jan et al., 2018; Jan et al., 2017a; Jan and Sigal, 2018; Jan et al., 2015; Jan et al., 2017b; Yang et al., 2018a; Yang et al., 2018b) PLM has proven extremely useful for the study of tissues ex vivo, revealing patterns of fiber architecture throughout the globes of humans and other animals. Importantly, the high angular resolution of PLM allows measurement of the degree of stretch or relaxation of collagen fibers, also referred to as crimp, that eludes other imaging techniques. Conventional PLM, however, requires multiple images acquired under various polarization states – four in our implementation. This slows down imaging, requires post-processing, such as image alignment, that affects image quality, and takes time. Hence, our studies using PLM were limited to static evaluation of tissues from eyes fixed at different intraocular pressures (IOPs). This was particularly limiting for the study of collagen fibers of lamina cribrosa (LC) beams, which vary so substantially even between contralateral eyes.

Recently we introduced instant polarized light microscopy (IPOL), a variation of PLM, which allows quantitative imaging of collagen at the full acquisition speed of the camera, with excellent spatial and angular resolutions. (Lee et al., 2019a; Yang et al., 2019) Our goal was to demonstrate IPOL for the quantitative analysis of both architecture and dynamics of ONH collagen. Specifically, we set out to visualize and quantify the microstructure and real-time dynamic response to stretch of collagen fibers of the peripapillary sclera (PPS) and LC. We give several examples that illustrate the great potential that dynamic visualization and quantification of the PPS and LC collagen with IPOL holds to help gain a better understanding of eye biomechanics and its role in health and disease.

## 2. Methods

This section is organized into three parts. First, we introduce IPOL imaging and how the technique produces true-color images indicating collagen fiber orientation. Second, we describe how to calibrate the color-angle mapping by imaging a section of chicken Achilles tendon at known orientations. Chicken tendon was chosen for its highly ordered and simple – compared with the eye – collagen fiber organization. (Jan et al., 2015; Yang et al., 2018b) Third, we utilized IPOL to image the ONH collagen architecture and illustrate the potential of the technique. We do this using fresh sheep eyes and a series of uniaxial and biaxial tests. Uniaxial stretch was applied to visualize continuous tissue deformations in a small region. Uniaxial stretch is a fairly common technique when evaluating the mechanical behavior of highly anisotropic materials, such as tendon. (York et al., 2014) Biaxial stretch was applied to visualize quasi-static deformations of the entire ONH region. Biaxial stretch is useful to reveal collagen fiber realignment, including rotation and torsion, and is a better approximation to the physiologic loading of scleral tissue. (Chung et al., 2016) From the IPOL images we analyzed stretch-induced collagen deformation and uncrimping in the PPS and LC, and changes in LC pore morphology as illustrative of the multitude of microarchitecture data available from IPOL. The rationale and implications of our choices of illustrative examples and testing techniques are addressed in more detail in the Discussion.

### 2.1 IPOL imaging system

IPOL was implemented with a commercial inverted microscope (IX83; Olympus, Tokyo, Japan), as previously described. (Lee et al., 2019b; Yang et al., 2019; Yang et al., 2021) Briefly, a broadband white-light source, a set of polarization encoder and decoder (each consisting of a polarizer and a polarization rotator), and a color camera (acA1920-155uc, Basler AG, Ahrensburg, Germany) were used in this system. In the absence of a birefringent sample, the white light was blocked and the image background appeared dark (Figure 1a). With a birefringent sample, such as collagen, the spectrum of the white light was changed and the light appeared colorful (Figure 1b). The colorful light was acquired by a color camera to produce the true-color images indicating collagen fiber orientation (Figure 1c). The frame rate of IPOL was limited only by that of the color camera, which in our setup was 156 frames per second. The high-speed imaging performance of IPOL allows capturing stretch-induced tissue deformation in real time, which is essential in characterizing tissue mechanics. (Ethier et al., 2004b)

**Figure 1.**
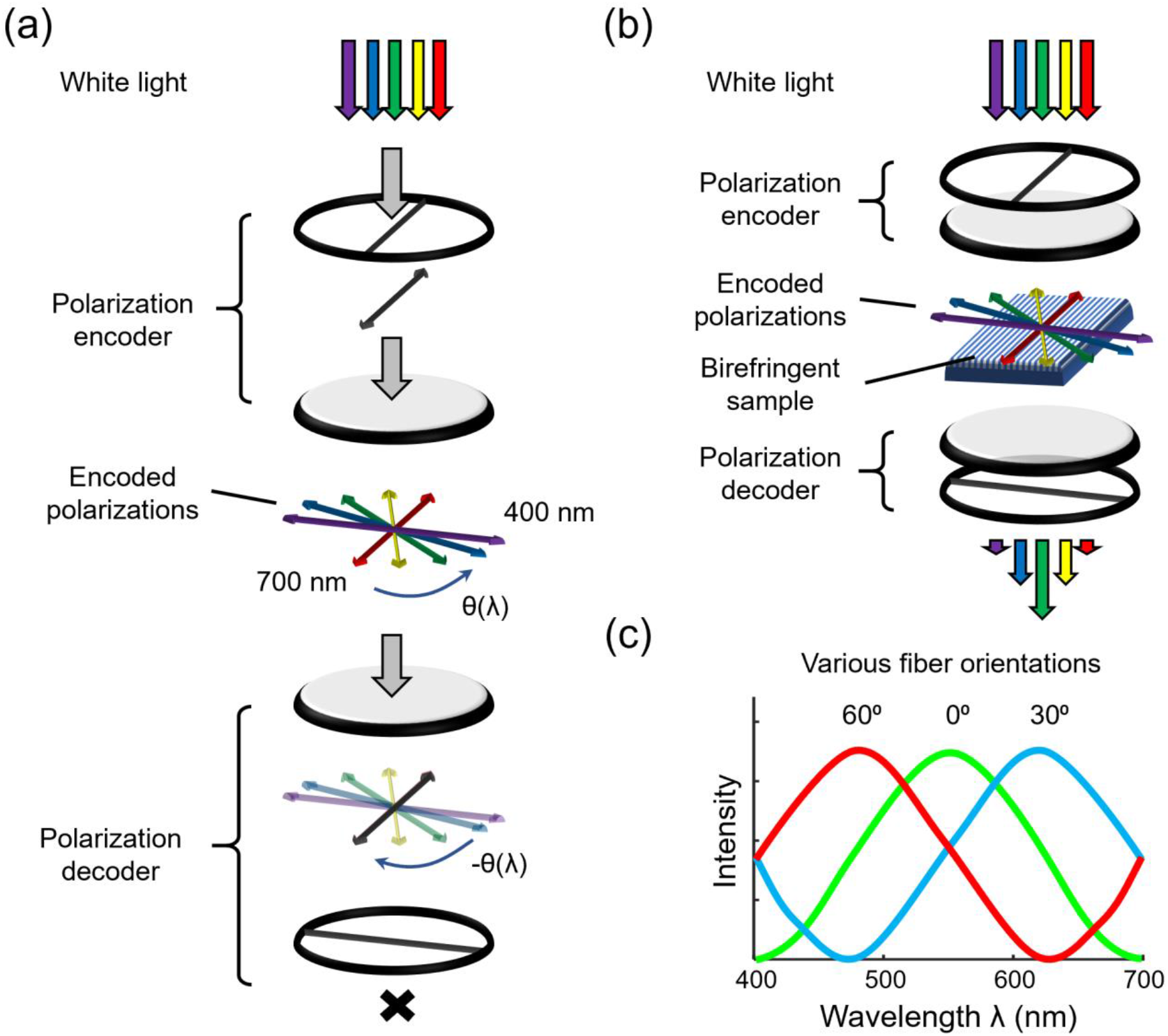
Schematic diagram of IPOL imaging system. **(a)** The white light passing through the polarization encoder was linearly polarized. The polarization directions of the spectrum were diverged within 90 degrees. In the absence of a birefringent sample, the polarization decoder did not allow light to pass through the system. **(b)** As the encoded polarized light passed through a birefringent sample, such as collagen, the aspect ratio of polarization of each wavelength was modulated based on the collagen fiber orientation. A new spectrum was then generated after the modulated polarized light passed through the polarization decoder. **(c)** The colorful light was acquired by a color camera to produce true-color images indicating collagen fiber orientation.

### 2.2 System calibration

#### Sample preparation

A chicken Achilles tendon was dissected and fixed with 10% formalin for 24 hours while under load to remove the natural undulations of collagen fibers, or crimp. (Yang et al., 2018b) Following fixation, the tendon was cryo-sectioned longitudinally into 30-µm-thick sections.

#### Color-angle mapping

IPOL images were acquired with the chicken tendon section at several controlled angles relative to the longitudinal fiber direction, from 0 to 90 degrees, every 2 degrees. The individual images were then registered using Fiji software. (Schindelin et al., 2012) A region of interest (ROI) on the tissue was manually selected on the stack, and the hue of the ROI was extracted by converting RGB (Red, Green, Blue) images into HSV (Hue, Saturation, Value) images. A color-angle conversion map was then computed by a circular interpolation of the measured hue and its corresponding orientation angle. The fiber orientation map for all images was then obtained by searching hue value over the color-angle conversion map for corresponding angles for all pixels (Figure 2).

**Figure 2.**
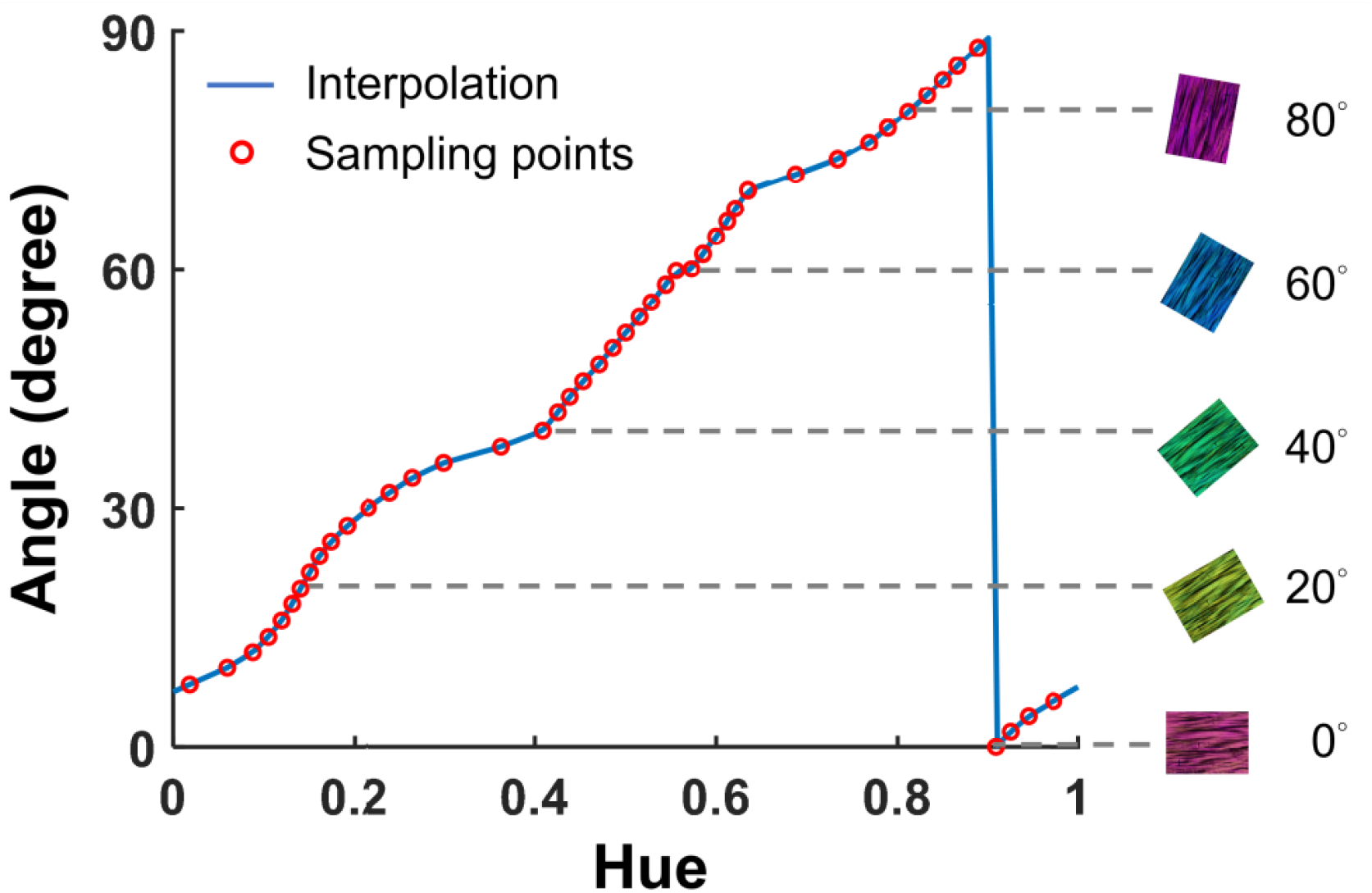
IPOL calibration curve. The curve was obtained from the circular interpolation of fiber orientation as a function of hue (*i*.*e*., the parameter of a color as determined by its dominant wavelength). Shown on the right are five example IPOL images of the same chicken tendon section at known orientations. Details of the calibration are described elsewhere. (Yang et al., 2021)

### 2.3 Imaging ONH collagen architecture and deformation

#### Sample preparation

Normal sheep eyes about a year old were procured from a local abattoir within four hours after death. The muscles, fat, and episcleral tissues were carefully removed. The ONH region was isolated using an 11-mm-diameter trephine and embedded in optimum cutting temperature (OCT) compound (Tissue-Plus; Fisher Healthcare, TX, USA). Samples were then snap frozen in liquid nitrogen-cooled isopentane and sectioned coronally at a thickness of 16 µm. OCT was washed with multiple PBS baths. To prevent curling or tears at the clamp points, a tissue section was sandwiched between two pieces of silicone sheet (Medical Grade, 0.005”; BioPlexus, AZ, USA). The sheets also allowed using PBS to maintain tissue hydration without lensing.

#### ONH collagen architecture

Tissue samples were imaged using a 4x objective (numerical aperture [NA], 0.13). Due to the limited field of view of the objective, the image of the whole section was acquired using mosaicking. The mosaics were obtained with 20% overlap and stitched using Fiji. (Schindelin et al., 2012) We have previously shown that the visualization of collagen fibers is not affected by mosaics or stitching. (Jan et al., 2018; Jan et al., 2015)

#### Uniaxial stretch testing

For uniaxial testing, sections were mounted to a commercial uniaxial stretching device using custom-made clamps (Microvice; S.T. Japan, FL, USA). The clamps allowed us to set the sample at the focal plane of our optical system, which was different from the commercial device default axis. The stretching process was imaged using IPOL in real-time display mode (156 frames per second) with a 10x strain-free objective (NA, 0.3). For each testing, either the LC or the PPS region was imaged. Our measurements were:

- *Maximum principal strain in the PPS*. Maximum principal strain was used to analyze the local deformation of the PPS. A digital image correlation technique was used to quantify the tissue displacement, and the maximum principal strain (a measure of tensile strain) was then calculated as described elsewhere. (Wei et al., 2018)
- Orientation profile *of LC beams*. The change in fiber orientation profile determines the crimp geometry of the collagen. The fiber orientation profiles along a beam were measured with stretch. The crimp tortuosity of LC beams was then calculated as previously described. (Brazile et al., 2018; Jan et al., 2018)

#### Biaxial stretch testing

Biaxial stretch testing is used to simultaneously extend the entire ONH region equally along two axes, which is a better mimic of the in vivo inflation conditions than uniaxial loading. Each section was mounted to a custom biaxial stretching device and then stretched quasi statically (small stretch steps [<0.1%] followed by long (20s or more) pauses to allow dissipating viscoelastic effects).(Lee et al., 2019a) At each step, multiple images were captured to cover the entire ONH region in a mosaic. The section was imaged using IPOL with a 4x strain-free objective (NA, 0.13). Our measurements were:

- *Angle distribution in the PPS*. Changes in crimp are central to the nonlinear behavior of the tissues. The changes in angle distribution along a bundle indicate if the loading causes bundle rotation or torsion. Angles within a PPS region were extracted at the initial and stretched states. We then used the Epanechnikov kernel to fit the angle distribution at each state. (Bowman and Azzalini, 1997)
- *LC beam width*. The test determines if the change in LC beam width under the stretch is related to crimp tortuosity of LC beams. We evaluated LC beam brightness profile across three beams and then estimated the beam widths using the full-width at half-maximum (FWHM). (Weik, 2001)
- *LC pore geometry*. Changes in LC pore geometry determines if LC pore deformation was isotropic. We traced 17 LC pores at the initial and stretched states, and then overlapped them to visualize pore deformation. Pore size (area) and shape (convexity and aspect ratio) were measured, (Voorhees et al., 2017a) and a paired *t*-test was used to evaluate their differences between the initial and stretched states. We used α = 0.05 to establish significance.

## 3. Results

### 3.1 ONH collagen architecture

Figure 3 shows an example IPOL image mosaic acquired of a static coronal section through the ONH of a fresh sheep eye. The image illustrates the high spatial and angular resolutions of IPOL, which allow identification of several aspects of collagen architecture simultaneously. At the smaller scale, crimp, or the natural waviness of collagen fibers, is discernible in both the PPS and LC regions. At the medium scale we can recognize the LC beams, and the width and orientation of fiber bundles. The larger scale we discern the overall shape of the scleral canal and the general organization of collagen in the PPS.

**Figure 3.**
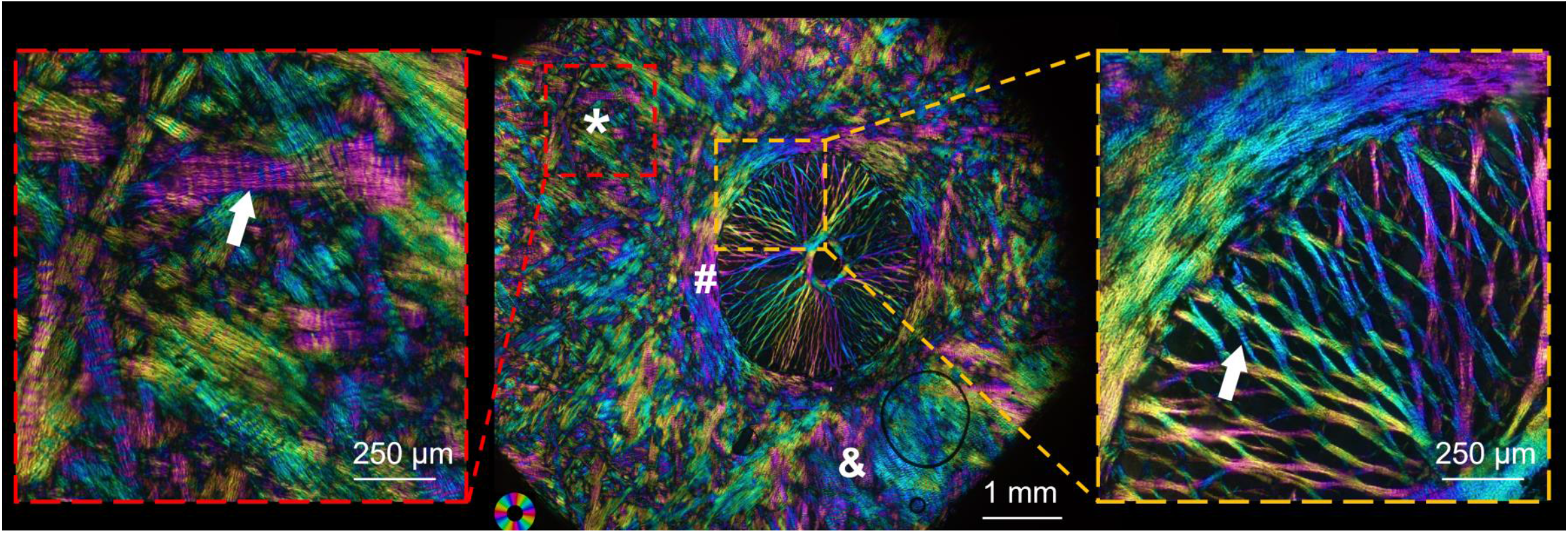
An IPOL image that was a mosaic acquired of a static coronal section from the ONH of a fresh sheep eye. Color in the image represents local fiber orientation and density. The black rings are optical artifacts due to air bubbles. In the PPS, there were fibers aligned circumferentially around the canal (#), fibers oriented radially from the canal (&), and unaligned interweaving fibers (*). In the LC, intersecting fibers formed large knots and wrapped around neural tissue pores, with beam insertions into the scleral canal wall that were either narrow and straight or wide and flared. Close-up shows fiber interweaving in the PPS (red box) and intersecting beams in the LC (yellow box). Crimp, or the natural waviness of collagen fibers is clearly discernible (marked by the white arrows; undulations in color).

### 3.2 Uniaxial stretch testing

Figure 4 illustrates local nonlinear deformations in a sample of PPS under uniaxial stretch visible using IPOL (Figure 4a). Tracking the fibers under stretch revealed that motion trails that were non-monotonic, inhomogeneous, tortuous and anisotropic (Figure 4b). A contour plot of the maximum principal strain shows that stretch-induced PPS deformation was non-affine, *i*.*e*., the local deformation differs from the applied stretch. (Figure 4c).

**Figure 4.**
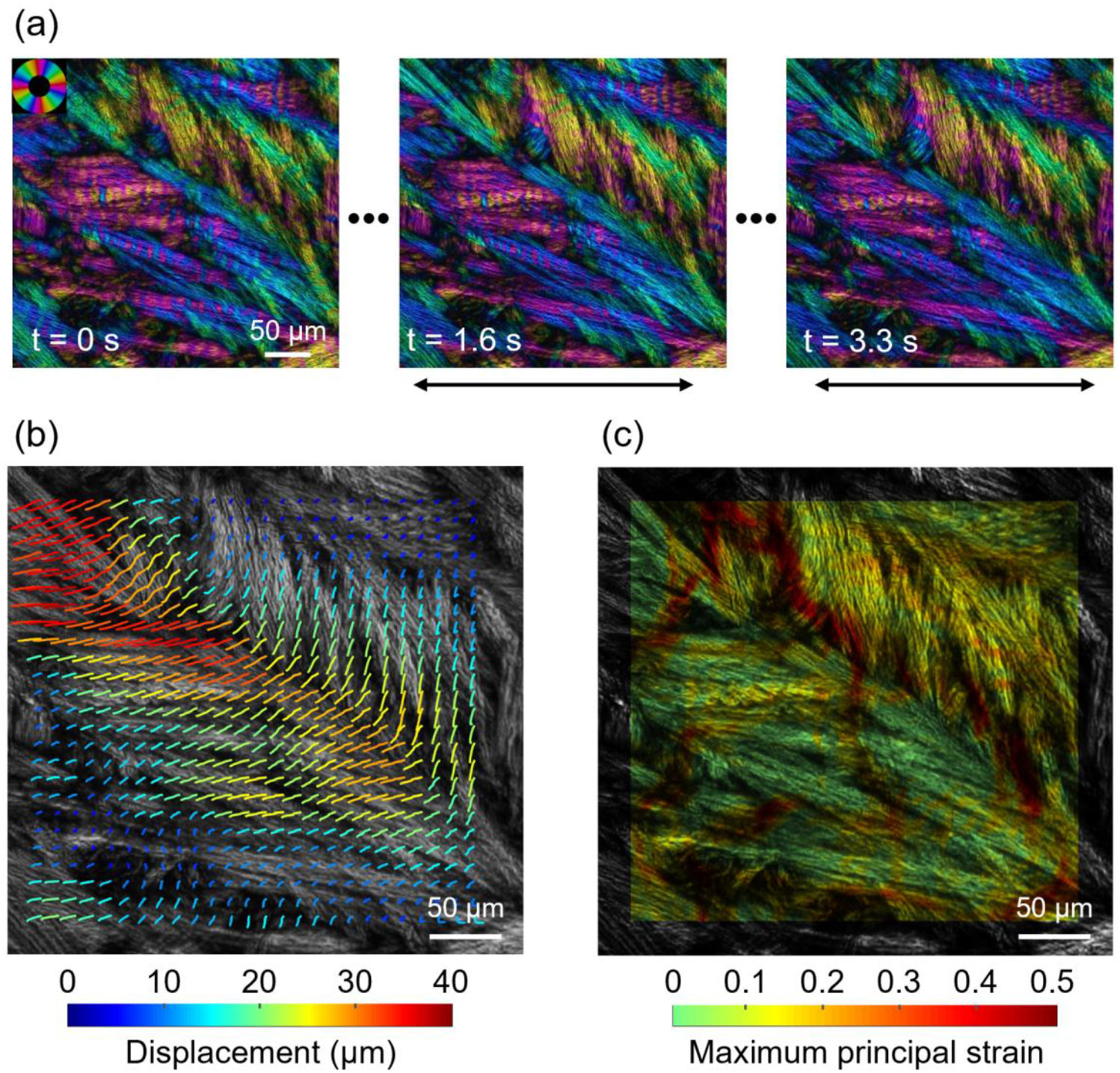
IPOL can capture highly dynamic deformations of ocular tissues in detail. **(a)** Time-sequence IPOL images of the PPS under uniaxial stretch. The black arrows indicate the stretch direction. 100 images were captured within 3.3 s. We show three images here to indicate the initial (t = 0 s), middle (t = 1.6 s), and final (t = 3.3 s) states. **(b)** Traces of stretch-induced tissue displacement. The motion trails were tortuous and anisotropic. For better visualization of tissue displacement, the color IPOL image was transformed into the grayscale image. **(c)** Contour plot of the maximum principal strain. Despite the PPS was under uniaxial stretch, its deformation was non-affine, *i*.*e*., the local deformation differs from the applied stretch. High strains were mainly located at fibers oriented transversely to the stretch direction. The peak strain reached 50%. It is important to note that these deformations were obtained in a well-controlled stretch test, which are likely different from in vivo or in situ, but that nevertheless provide a valuable opportunity to study the inter-relationship of structure and mechanics in ocular tissues.

The uncrimping process of an LC beam under uniaxial stretch is shown in Figure 5. Before stretch, a beam exhibits collagen fiber undulations (Figure 5a t = 0 ms). The undulations are discernible both based on the fiber edges and by the color bands. The crimp bands are perpendicular to the principal beam axis. Elsewhere we had speculated that these properties would cause the beam to stretch without torsion.(Jan et al., 2017a) As the beam was stretched, the undulations decreased, according to the expected collagen uncrimping (Figure 5a, t = 400 ms to t = 1200 ms). Taking advantage of IPOL quantitative information on the local fiber orientation, we plotted the orientations along a line along the beam axis (Figure 5b). The plot illustrates the regular sinusoidal nature of the crimp. The amplitude of the fiber orientation profile along the beam decreased as the stretch increased, another indication of collagen uncrimping. Interestingly, the uncrimping is not associated with a discernible increase in crimp period. This is consistent with our measurements in sclera and can be understood given the small undulation angles. (Jan and Sigal, 2018) Interestingly, the largest undulations uncrimped earlier in the stretch, such that the crimp was smaller and more uniform after stretch than before stretch. The uncrimping process can be further quantified by calculating the tortuosity as a function of time, which can potentially be used as input for constitutive model (Figure 5c). (Grytz and Meschke, 2009; Hill et al., 2012)

**Figure 5.**
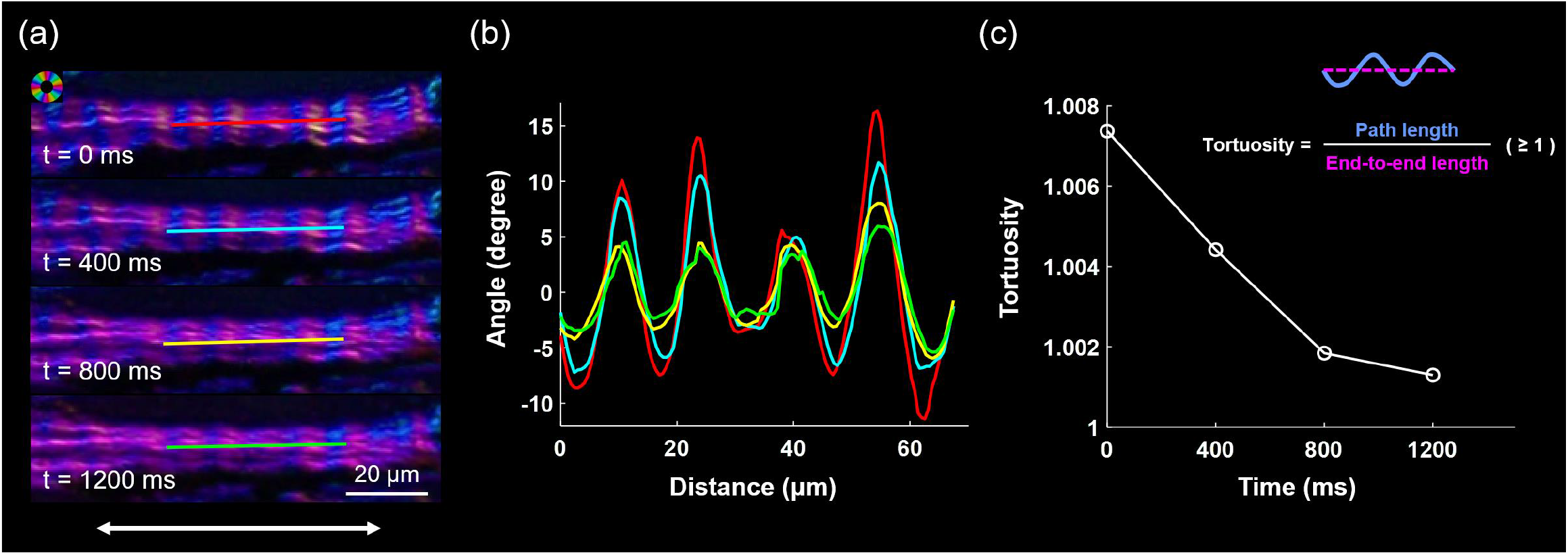
Uncrimping process of an LC beam under uniaxial stretch. **(a)** IPOL images of an LC beam captured at four time points with an interval of 400 ms. The white arrows indicate the stretch direction. Before stretch (t = 0 ms), the beam was crimped, as evidenced by color undulations along the beam, without torsion. As the stretch increased, there were less undulations in color, suggesting a reduction of crimp. **(b)** Fiber orientation profiles along the solid lines in (a), with colors corresponding to different time points. Before stretch, the orientation profile (red curve) exhibited periodicity with angles alternating between −10 and 15 degrees. As the stretch increased, the amplitude of the orientation profile decreased, indicating collagen uncrimping. Based on the difference in the period of the orientation profiles between the initial (red curve) and final (green curve) states, the stretch of the LC beam was measured as 3.6%. At this stretch level, the LC beam did not reach full uncrimping. **(c)** The high spatial and angular details allow measurement of crimp tortuosity along the beam. As expected, the tortuosity or degree of undulation, decreases with time and stretch.

### 3.3 Biaxial stretch testing

Deformations of the PPS and LC under biaxial stretch are shown in Figure 6. In the PPS, the angle distribution changed from bimodal at the initial state, indicating crimped fibers, into unimodal at the stretched state (Figure 6a). The peak angle after stretch is not within the two initial peaks, indicating that the uncrimping also involved a rotation of the fiber bundle. In the LC, crimp tortuosity and beam decreased under the stretch (Figure 6b). However, we also noticed something interesting worth pointing out: the beam with the least crimped collagen at the initial state narrowed the most at the stretched state.

**Figure 6.**
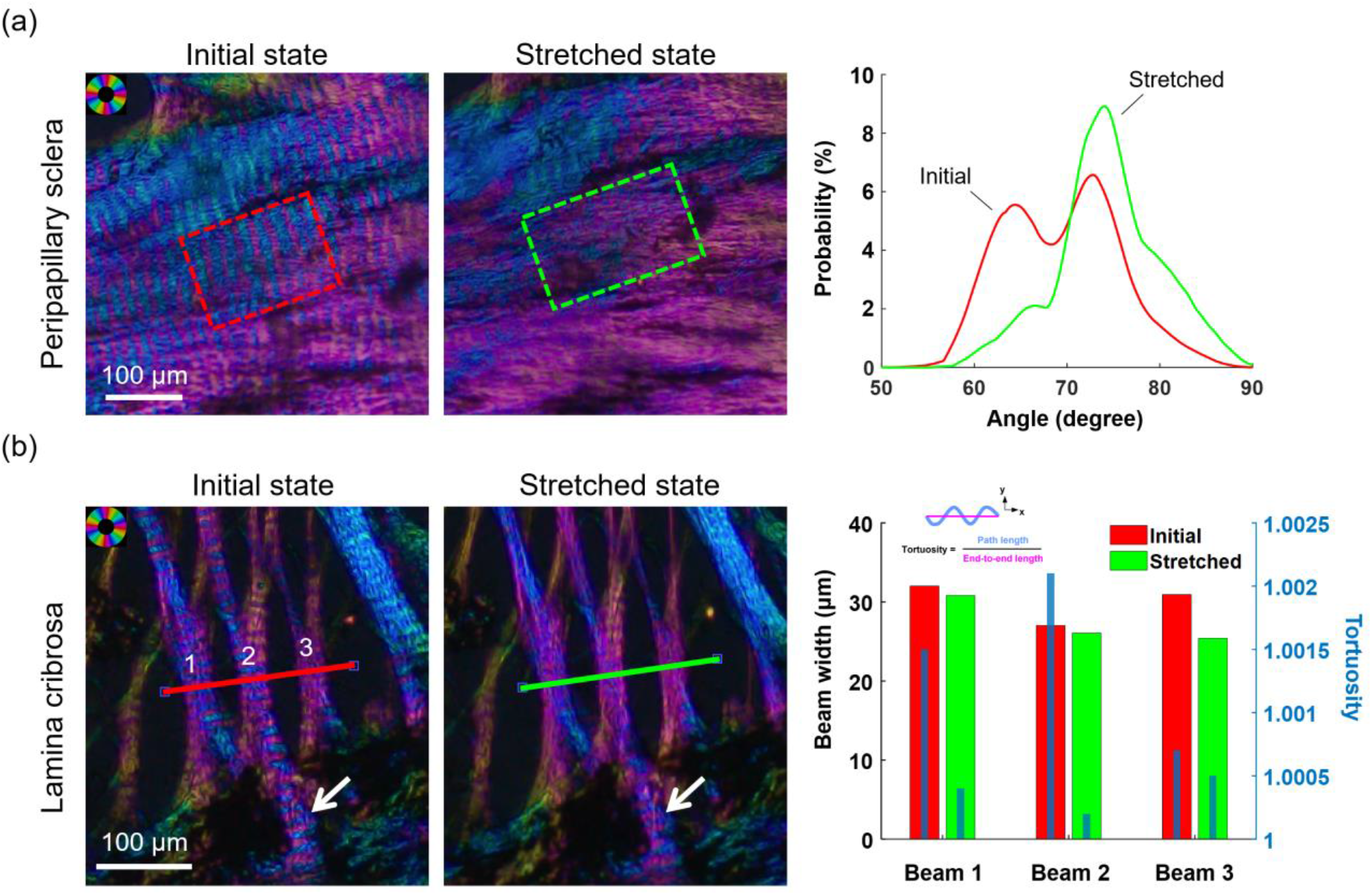
PPS and LC fiber uncrimping under biaxial stretch. **(a)** IPOL images of the PPS at the initial and stretched states. To quantify stretch-induced changes in collagen crimp, angle distributions were measured within a region of interest (ROI, dashed rectangle) and compared before and after stretch. At the initial state, the angle distribution within the ROI exhibited a bimodal pattern, indicating crimped fibers. In the stretched state, the distribution changed into a unimodal pattern due to uncrimping. **(b)** IPOL images of the LC at the initial and stretched states. We measured the crimp tortuosity and width (along the dashed line) of three beams. Before stretch, the tortuosity of Beams 1, 2, and 3 was 1.0032, 1.0048, and 1.0011, and their width was 32, 27, and 31 μm, respectively. After stretch, both the tortuosity and width of the three beams decreased. The tortuosity decreased by 72%, 69%, and 46%, and the width decreased by 3.8%, 3.5%, and 18%, respectively. Beam 3, which was less crimped (*i*.*e*., smaller tortuosity) than Beams 1 and 2 before stretch, narrowed more after stretch.

Figure 7 shows an example of analysis of stretch-related LC biomechanics at the pore level. Comparing pores before and after stretch revealed that stretch-induced pore deformation was anisotropic within a single pore and non-uniform among pores. Pores became larger, more convex, and more circular after stretch (*P*’s < 0.005).

**Figure 7.**
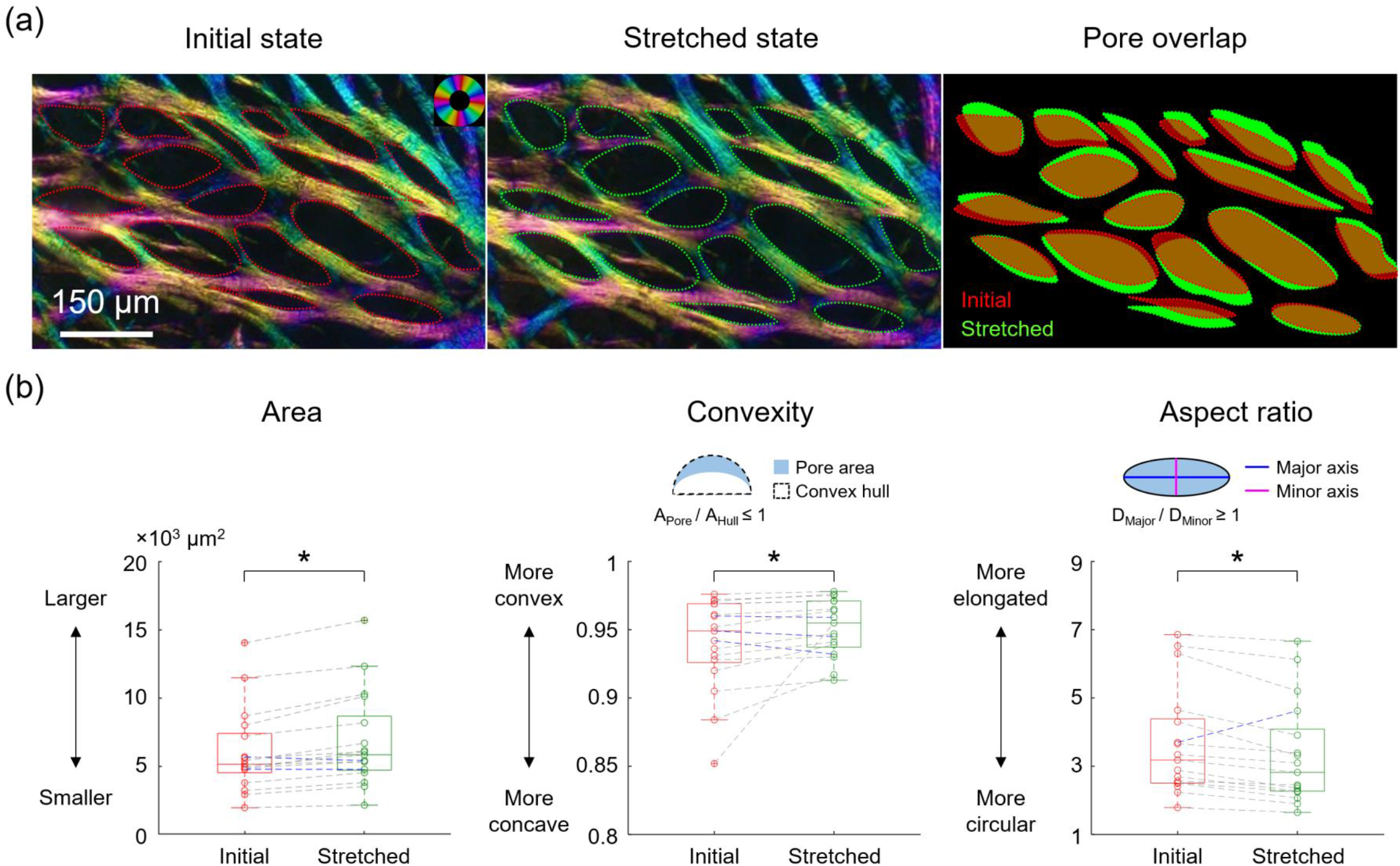
Deformation of LC pores under biaxial stretch. **(a)** 17 pores were traced at the initial and stretched states, and then overlaid to show the differences in pore contour shape and size before and after stretch. The overlapped image shows that stretch-induced pore deformation was anisotropic within a single pore and non-uniform among pores, indicating that neural tissues and vessels within pores suffer shear deformation. **(b)** Box plots of area, convexity, and aspect ratio of the 17 pores at the initial and stretched states. The dotted lines in the box plots connect the data points corresponding to the same pore at the initial and stretched states, where the blue dotted lines represent the opposite trend against the majority of the pore changes. After stretch, 15 and 14 pores have increases in pore area and convexity, respectively, and 16 pores have a decrease in pore aspect ratio; the average pore area and convexity increased by 13% and 1.3%, respectively, and the average pore aspect ratio decreased by 9%. This suggests that pores became larger, more convex, and more circular after stretch. Convexity was defined as the ratio of the area of the pore to the area of the convex hull of the pore. Aspect ratio was defined as the ratio of the major axis length to the minor axis length. The superscript * indicates *P* < 0.05.

## 4. Discussion

Our results demonstrate that IPOL is well-suited for the imaging and quantitative study of the architecture and dynamics of ONH collagen. Using IPOL we made several interesting observations. First, stretch-induced PPS deformation was non-affine, non-linear and non-uniform. Second, the crimped collagen fibers in the PPS and LC straightened under stretch, without torsion. Third, stretch-induced LC pore deformation was anisotropic within a single pore and non-uniform among pores. Below we discuss each of these observations and their implications in ONH biomechanics.

We observed that the distribution of strains in the PPS under uniaxial stretch was non-uniform. The highest strains were mainly located at fiber bundles oriented transversely to the stretch direction. Compared to bundles along the stretch direction, transverse bundles are less efficient at carrying the loads, and thus experienced higher strains perpendicular to the fiber orientations. In addition, the stretch-induced PPS deformation was non-affine. This raises the possibility of complex modes of energy storage and dissipation. (Billiar and Sacks, 1997) A common assumption of numerical models of the PPS, and many other soft tissues, is that the microstructure undergoes affine deformations. (Coudrillier et al., 2013; Girard et al., 2009; Grytz et al., 2012; Voorhees et al., 2018) If our observations under uniaxial stretch carryover to the physiologic conditions, it would indicate a fundamental limitation of current models and the need for more complex models that can account for non-affine fiber behavior. (Wang et al., 2020)

We observed that the crimped collagen fibers in the PPS and LC straightened under stretch. This phenomenon is well recognized in other tissues, like tendon and ligament, (Hansen et al., 2002; Thornton et al., 2002) We have also imaged and characterized collagen crimp and recruitment in ONH tissues, but the PLM techniques available required us to do it in samples fixed under different conditions. (Jan and Sigal, 2018) To the best of our knowledge, this study is the first showing the uncrimping process of ONH collagen under stretch. The process we report is consistent with theory of collagen fiber recruitment in soft tissues, in which fibers straighten with stretch to become straight are considered recruited, contributing to the local stiffening of the tissue. (Grytz and Meschke, 2009; Hill et al., 2012; Holzapfel, 2001) Collagen uncrimping and recruitment is an essential mechanism in the mechanical response of the eye tissues to mechanical load and deformation. Hence, the ability to visualize and quantify crimp under dynamic tests that IPOL provides represents a powerful opportunity to characterize and understand the tissues, and how the tissue behavior arises from collagen microarchitecture and overall morphology. (Eilaghi et al., 2010; Girard et al., 2009; Jan and Sigal, 2018; Perez et al., 2014)

Our results suggest that stretch-induced changes in an LC beam width may be related to its microstructural crimp characteristics. Specifically, less crimped LC beams narrowed more under stretch (a Poisson effect). This may affect the adjacent neural tissues due to the lateral movement of the beam edges, and may cause increased stress within the beam due to decreased cross-sectional area. Further work is needed to examine the spatial relationship between collagen crimp in the LC and stretch-induced LC beam narrowing.

Analyzing stretch-induced LC pore deformation is critical to understand the biomechanical insult to the neural tissues and vessels within the pores resulting from IOP. (Ling et al., 2019; Sigal et al., 2007; Sigal et al., 2014a; Voorhees et al., 2017a; Voorhees et al., 2017b) Using IPOL, we observed that stretch-induced LC pore deformation was anisotropic within a single pore and non-uniform among pores. This indicates that neural tissues and vessels within LC pores may experience multiple modes of deformation under stretch, such as stretch, compression, and shear. Quantitative analysis further shows that pores became larger, more convex, and more circular after stretch. This is consistent with numerical findings of IOP-induced changes in LC pore morphology. (Voorhees et al., 2017a) Understanding how stretch influences pore deformation will help clarify the association between the distribution of lamina pore shape and glaucoma status or progress, which remains controversial in the literature. (Akagi et al., 2012; Fontana et al., 1998; Miller and Quigley, 1988; Wang et al., 2013; Zwillinger et al., 2016) Of course, it is essential to consider the conditions in which we did the experiments and the limitations these imply. We discuss these further down.

The IPOL imaging technique, in which a single snapshot image encodes information of fiber orientation and density, is uniquely well-suited to understanding the ONH collagen architecture and behavior under dynamic loading. IPOL provides high spatial and angular resolutions, which allows identifying collagen architectural details, such as fiber interweaving and crimp. Although conventional PLM can also identify these details, its resolution is lower (Figure 8). PLM requires post-processing (*i*.*e*., image registration and denoising) multiple images to derive information such as fiber orientation, which reduces the spatial detail of the image. IPOL also has high temporal resolution, limited only by the frame rate of the camera. Hence, IPOL is suited for imaging continuous tissue deformation under dynamic loading. Conventional PLM can only be employed to image quasi-static tissue deformation, in which the dynamic process has to be split into multiple static steps. (Tower et al., 2002) There are alternative faster PLM techniques, for example using multiplexed analyzer filters, but these reduce the resolution. (Gruev et al., 2010)

**Figure 8.**
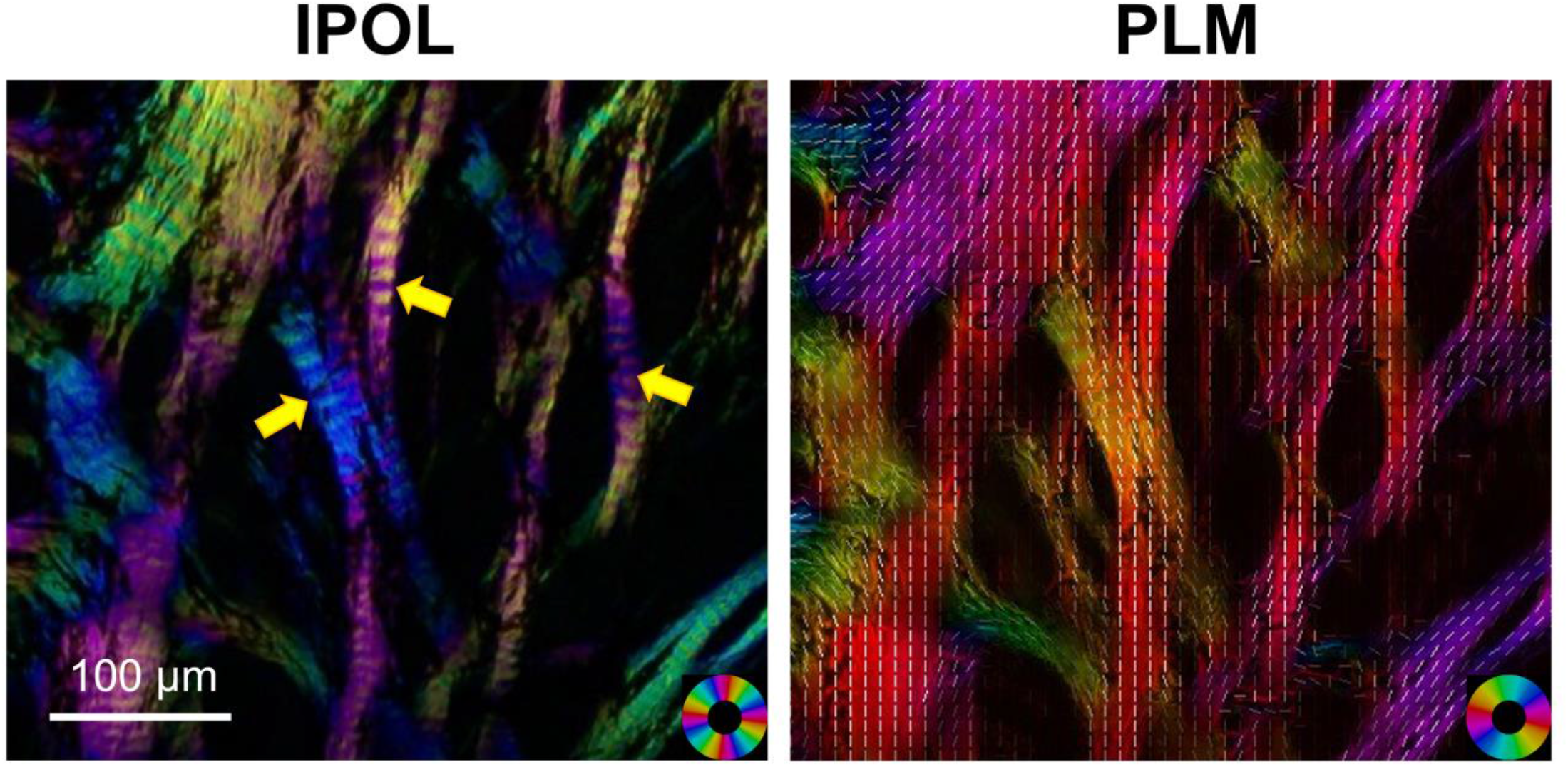
A comparison between the IPOL (left) and PLM (right) images of a sheep ONH section. The collagen crimp of the LC beams (yellow arrows) is easily discernible in the IPOL image due to its high spatial and angular resolutions. Note that the orientation-encoded colors repeat every 90 degrees in IPOL and every 180 degrees in PLM. The white lines in the PLM image represent local fiber orientations.

It is important to consider the limitations of IPOL and our analysis in this work. First, we used sheep eyes to study the ONH collagen deformation under stretch. Sheep eyes are similar to human eyes in that they have a collagenous LC, but differ in having a ventral groove in the ONH, similar to that in pig. (Brooks et al., 1998) Though it is possible that stretch-induced ONH collagen deformation found in sheep is not the same in humans, it is important to understand sheep as an animal model. (Candia et al., 2014; Gerometta et al., 2010) Future work should include additional animal models as well as human eyes. The sheep from which we obtained the eyes were young, as is to be expected from an abattoir. We have shown that crimp in the eye decreases with age, (Gogola et al., 2018a) and thus it is possible that older eyes behave differently.

Second, IPOL is based on transmitted light illumination, which requires the tissue samples to be cut into fairly thin sections (thickness of 16 µm in this study). Thus, while the architecture of collagen fibers in the samples is three-dimensional, we have limited the sample to a two-dimensional “slab”. Although some publications have obtained valuable information on ONH biomechanics by focusing on the plane behavior, (Voorhees et al., 2017a; Voorhees et al., 2017b) this is a major simplification that likely has a substantial effect on stretch-induced ONH collagen deformations. The approach used herein, however, does have one important positive aspect: the collagen visible in the images is all there is in the sample. This is in contrast to the vast majority of experiments of the posterior pole in which only part of the sample is visible. For example, in inflation experiments it may be possible to measure deformations of the sclera surface, but any mechanics within the tissues must be assumed or inferred. (Bruno et al., 2018; Geraghty et al., 2020)

Third, ONH sections were stretched either uniaxially or biaxially. This loading condition is a fairly common assumption in studies on ONH biomechanics when the intent is to simulate the tension in the ONH induced by IOP. (Perez et al., 2014; Voorhees et al., 2017b; Zhang et al., 2015) It remains to be determined whether the ONH collagen deformation identified in this study extends to physiological loading conditions or not. Also, whilst it is fairly likely that the loading applied to the scleral samples is not physiologic, the conditions may be more realistic for LC beams. Because the neural tissues adjacent to the beams are substantially more compliant than the beam, the assumptions of in-plane stretch along the beam may approximate the physiological condition better.

Fourth, the orientation-encoded colors in IPOL are cyclic, *i*.*e*., repeating every 90 degrees. As a result, two fibers oriented perpendicularly to each other exhibit the same color. Based on color-angle mapping, these two fibers would have the same quantified orientation angles. This may affect the quantification of collagen fiber interweaving and large rotation of collagen fibers under stretch, which was beyond the scope of this study. It is possible to modify the setup to create a system repeating every 180 degrees. This, however, may adversely affect angular resolution.

Fifth, to capture the deformation of the entire ONH region under biaxial stretch, we stitched multiple images into mosaics at each stretch level. Mosaicking is only suitable for imaging quasi-static tissue deformation and may cause image artifacts such as imperfect registration, poor or irregular focus, or inconsistent illumination. Implementing IPOL with a dissecting microscope with a broader depth of field and a deeper focal depth can achieve a large field of view of a single image and thus circumvents the need for mosaicking, potentially at the cost of resolving power. (Lee et al., 2020)

In conclusion, we have demonstrated that IPOL allows visualization and quantification of changes in ONH collagen morphology and architecture under dynamic loading. This study represents an important step towards using a novel imaging modality to study ONH collagen microstructure and biomechanics, which could help understand the role of collagen microstructure in eye physiology, aging, and in biomechanics-related diseases, such as glaucoma and myopia.

## Acknowledgements

This work was supported in part by National Institutes of Health grants R01-EY023966, R01-EY028662, P30-EY008098 and T32-EY017271 (Bethesda, MD), and the Eye and Ear Foundation (Pittsburgh, PA), Research to Prevent Blindness (New York, NY).

## Notes

**Funding:** Supported in part by National Institutes of Health R01-EY023966, R01-EY028662, P30- EY008098 and T32-EY017271 (Bethesda, MD), the Eye and Ear Foundation (Pittsburgh, PA), and Research to prevent blindness.

### Competing Interest Statement

Ziyi Zhu was at the University of Pittsburgh when he contributed to this work. He is now at Amazon; Other authors have nothing to disclose.

